# A flexible modelling framework for estimating thermal tolerance and sensitivity

**DOI:** 10.64898/2026.07.16.738378

**Authors:** Daniel W.A. Noble, Pieter A. Arnold, Shinichi Nakagawa, Patrice Pottier

## Abstract

Extreme heat events are becoming more frequent, intense and prolonged, making it urgent to predict how heat intensity and exposure duration combine to threaten organisms. Thermal death time (TDT) and thermal load sensitivity (TLS) models provide this link, but conventional two-stage analyses often discard uncertainty, mishandle censored or overdispersed data and limit inference. Here, we show how the four-parameter log-logistic model can recover TDT/TLS quantities, including thermal tolerance (*CT*_*max*_), sensitivity (*z*), critical temperature (*T*_*crit*_), heat injury and survival, from one model. Simulations show the joint model reproduces classical estimates when two-stage assumptions hold and is more reliable when they fail. We introduce these workflows as Bayesian and frequentist R packages. Case studies across plant and animal taxa demonstrate this modelling framework can estimate group contrasts and predict survival from realistic field temperature-time series. This framework provides more robust inference and flexible tools for predicting organismal responses to extreme heat events.

## Introduction

Climate change is increasing the frequency, intensity, and duration of extreme heat events globally, with cascading consequences for organismal fitness, population persistence, and biodiversity (Arnold *et al*. 2025; Rezende *et al*. 2020; Smale *et al*. 2019; Snook *et al*. 2026; Wiens 2016). Quantifying variation in thermal tolerance within and among species is central to forecasting biological responses to climate change in nature (Comte & Olden 2017; Deutsch *et al*. 2008; Duarte *et al*. 2012; Jørgensen *et al*. 2022; Pinsky *et al*. 2019; Sunday *et al*. 2011, 2012). Threshold-based measures of heat tolerance, such as the critical thermal maximum (*CT*_*max*_), have long been used to understand differences in sensitivity among taxa and to predict vulnerability to climate change (Deutsch *et al*. 2008; Pinsky *et al*. 2019; Pottier *et al*. 2025; Sunday *et al*. 2011, 2012). While useful for comparisons, threshold-based metrics do not capture how the biological effects of heat stress on fitness depend on both exposure duration and intensity, limiting our ability to make realistic predictions in nature (Clusella-Trullas *et al*. 2021; Jørgensen *et al*. 2021; Ørsted *et al*. 2022; Rezende *et al*. 2014; Troia 2023).

The Thermal Death Time (TDT) and Thermal Load Sensitivity (TLS) frameworks address this limitation by quantifying how the intensity and duration of heat exposure jointly determine survival and other fitness-related responses (Arnold *et al*. 2025; Cook *et al*. 2024; Jørgensen *et al*. 2019, 2022; Ørsted *et al*. 2022; Rezende *et al*. 2014). Recent extensions apply the same logic to sublethal endpoints, such as sterility and heat coma (Arnold *et al*. 2025; Ørsted *et al*. 2024; van Heerwaarden & Sgrò 2021). These models describe heat injury as a dose-dependent process above a critical threshold (*T*_*crit*_). Thermal injury accumulates more rapidly at higher temperatures and during longer exposures, and organisms may eventually reach a thermal load that compromises growth, reproduction, or survival if injury cannot be repaired (Arnold *et al*. 2025; Buckley *et al*. 2025; Jørgensen *et al*. 2019, 2021; Ørsted *et al*. 2022; Rezende *et al*. 2014). By linking biological responses to the intensity and duration of heat stress, TDT/TLS frameworks allow more realistic predictions under fluctuating thermal regimes (Jørgensen *et al*. 2021; Ørsted *et al*. 2022; Rezende *et al*. 2014), and have shown promise in predicting observed mortality rates in nature (Molina *et al*. 2025; Rezende *et al*. 2020; Verberk *et al*. 2023; Zamora *et al*. 2026).

Despite their potential predictive power, TDT/TLS analyses traditionally involve a two-stage modelling approach (Jørgensen *et al*. 2019; Li *et al*. 2023; Ørsted *et al*. 2022; Rezende *et al*. 2020; Truebano *et al*. 2018). First, organisms are exposed to multiple temperatures for different durations, and the proportion of individuals that are dead, infertile, knocked down, or otherwise impaired or affected is modelled as a function of exposure duration. This is typically done by fitting independent dose-response curves at each assay temperature (Figure 1a). From these curves, the exposure duration required to reach a specified response threshold is estimated, most often *t*_50_, the time required to reach 50% mortality, infertility, knockdown, or some other endpoint. Second, the resulting log_10_(*t*_50_) estimates are regressed against assay temperature using linear regression. Thermal sensitivity (*z*), the temperature increase required to reduce *t*_50_ by a factor of 10, and critical thermal maxima 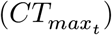 are then derived from this linear model (Ørsted *et al*. 2022; Rezende *et al*. 2014) (Figure 1b). Because 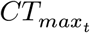 is estimated at a fixed exposure duration, it corresponds to the static critical thermal maximum, often denoted *sCT*_*max*_, in TDT/TLS studies (Jørgensen *et al*. 2019; Ørsted *et al*. 2022). We use *CT*_*max*_ throughout for brevity. 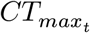 is usually computed for a 1-min (Rezende *et al*. 2014), 1-hour (Jørgensen *et al*. 2019), or 4-hour (Ørsted *et al*. 2024) duration of heat stress exposure. These quantities can then be used to estimate cumulative Heat Injury (HI) under natural temperatures, where injury accumulates as a function of the exposure duration relative to the predicted tolerance time at each temperature (Jørgensen *et al*. 2019; Villeneuve & White 2024). Recent extensions have emphasized that injury accumulation may also be offset by repair or acclimation mechanisms, motivating the emergence of new models that incorporate damage-repair dynamics, although applying such approaches remains challenging (Arnold *et al*. 2025; Buckley *et al*. 2025; Ørsted *et al*. 2026; Schow-Madsen *et al*. 2025; Wehrli *et al*. 2024).

**Figure 1:**
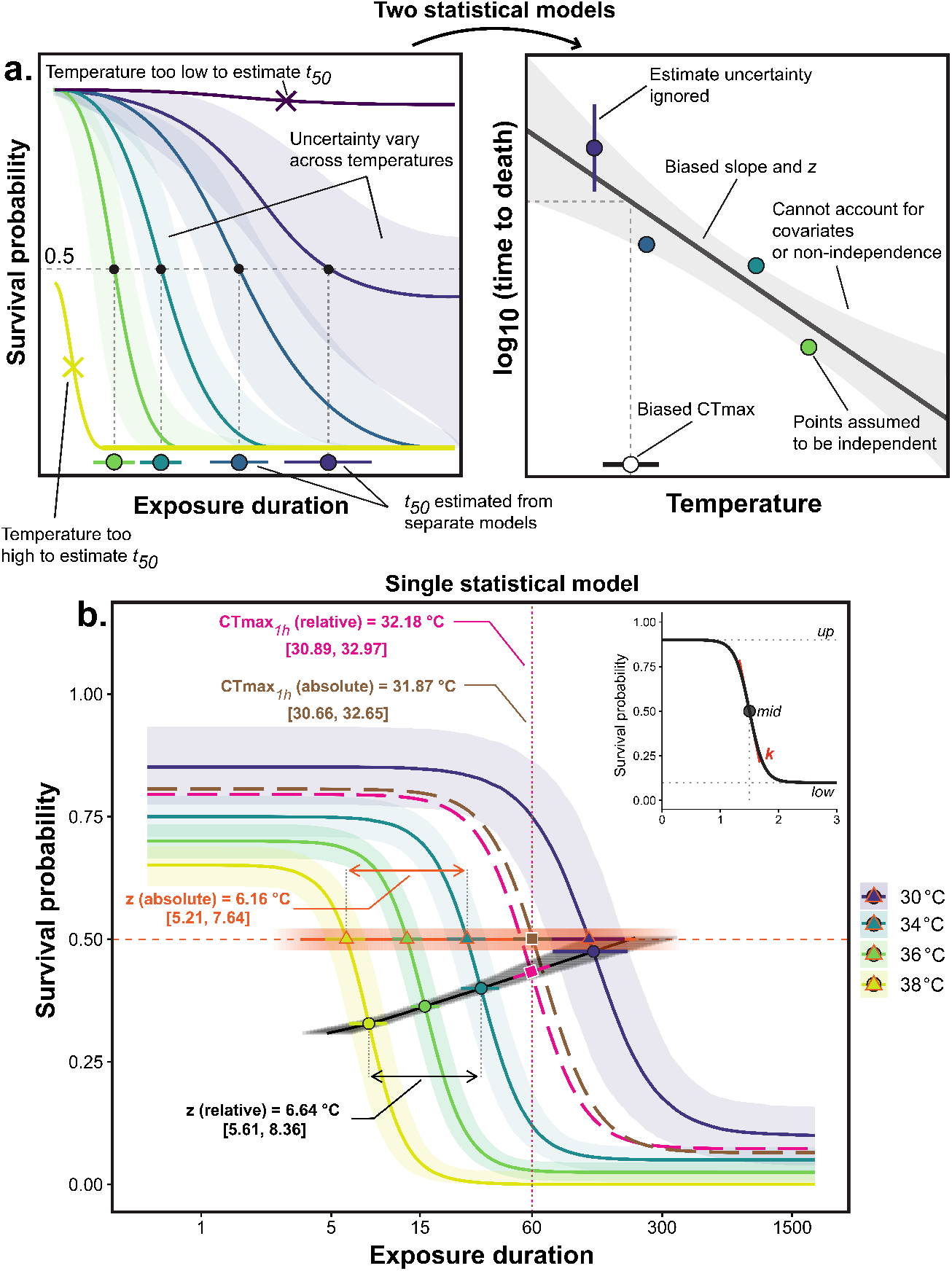
Conceptual comparison of the classical two-stage TDT/TLS workflow and the joint four-parameter log-logistic (4PL) model. (a) In the two-stage workflow, separate dose-response curves are fit at each assay temperature and the resulting *t*_50_ estimates are regressed against temperature. This workflow can fail when assays do not bracket *t*_50_, treats first-stage estimates as fixed and independent, and does not naturally propagate uncertainty or account for covariates / hierarchical structure into *z* and 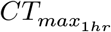. (b) The joint 4PL model estimates the full survival surface in one model. Coloured curves show posterior median survival probabilities at assay temperatures, with shaded pointwise 95% credible intervals. Filled circles mark the model midpoint, `mid(T)`, defining relative log_10_ *t*_50_; open triangles mark the absolute 50% survival crossing. A *relative* threshold is each curve’s own midpoint (halfway between its lower and upper asymptotes), whereas an *absolute* threshold is a fixed survival probability (here 50%); the two coincide only when the asymptotes are 0 and 1 (see Methods). The black and steelblue trajectories give the relative and absolute *t*_50_ relationships used to derive *z*_rel_, *z*_abs_, and 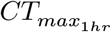 at the 60-min reference. Dashed curves show the fitted dose response at the corresponding relative and absolute 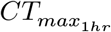 values. Inset: the 4PL parameters `low`, `up`, `mid`, and `k`. Values are illustrative.

While the two-stage pipeline is intuitive and easy to implement, it has several practical and statistical limitations (for a recent discussion of some issues, see Bullard *et al*. 2026). First, it rarely propagates uncertainty. The *t*_50_ estimates derived in the first stage have non-negligible sampling variance, but they are commonly treated as fixed-point estimates in the second-stage regression. Neglecting this sampling variance can bias or miscalibrate inference on *z* and *CT*_*max*_, and lead to inflated *R*^2^ values. Second, models estimating *t*_50_ often assume that upper and lower survival are 1 and 0, respectively. When data are sparse or do not span the response range, *t*_50_ estimates can be biased or undefined, leading to inaccurate *z* and *CT*_*max*_ estimates. Third, group comparisons between species, sexes, or life stages inherit miscalibrated uncertainty (as discussed by Lappi & Luoranen 2018), which affects inferences. Reducing each dose-response curve to a fixed *t*_50_ estimate also makes it difficult to determine which aspects of the dose-response curve are affected. Fourth, fitting separate dose-response models for each assay temperature assumes that these estimates are independent, and typically ignores the hierarchical structure of experimental data. Ignoring this nesting structure prevents the decomposition of variance and can lead to pseudoreplication, particularly when replicate-level variation affects the shape or position of the dose-response curve (Bullard *et al*. 2026). Finally, the two-stage pipeline does not easily allow downstream predictions of heat injury and survival in nature with uncertainty, which is key to inferring confidence in predictions of extreme-heat impacts.

Here, we present a single-stage alternative to the classical two-stage pipeline: a hierarchical four-parameter log-logistic model fitted jointly to all temperature-by-duration response data. We show how classical TDT/TLS quantities, such as thermal sensitivity (*z*), thermal tolerance (*CT*_*max*_), critical temperature thresholds (*T*_*crit*_), heat injury accumulation, survival under fluctuating thermal regimes, and their uncertainty, can be derived from this one model. We develop these tools as two R packages, `bayesTLS` (Bayesian) and `freqTLS` (frequentist), to facilitate their use across the community. We use simulations and empirical case studies across plant and animal taxa to demonstrate how to implement, validate, and extend this modelling framework.

### A non-linear hierarchical model for characterizing thermal load sensitivity

### Model specification

Assuming we have mortality data at different times and temperatures, let *y* denote the number of survivors among *n* individuals exposed for time *t* (e.g., minutes) at assay temperature *T* (°C). We can model the counts directly with a binomial (or beta-binomial) likelihood:

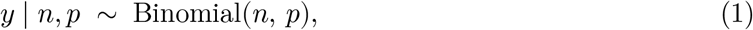

with survival probability *p* given by a four-parameter log-logistic (4PL) function of log_10_-exposure time:

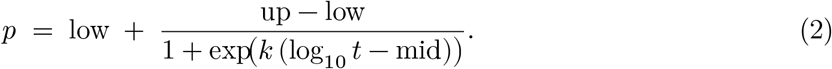

where low is the lower survival probability asymptote, up is the upper survival probability asymptote, and *k* is the steepness parameter controlling how rapidly survival probability declines on the log_10_ *t* axis (the slope of *p* versus log_10_ *t* at the midpoint is −*k*(up − low)/4). Larger *k* gives a sharper decline in survival probability with log_10_ *t*. mid is the value of log_10_ *t* at which *p* falls to the midpoint between low and up. The four-parameter log-logistic model provides a flexible sigmoidal description of thermal dose responses, with interpretable parameters for the lower asymptote, upper asymptote, curve steepness, and midpoint (Nielsen *et al*. 2004; Ritz 2010; Ritz *et al*. 2015; Ritz & Streibig 2008; Rudemo *et al*. 1989). We focus on binomial and beta-binomial outcomes because count data dominate TDT/TLS studies, but the same modelling logic can be extended to other response types using appropriate likelihoods, including continuous performance traits or latency data.

### Deriving key thermal load sensitivity parameters

The TDT/TLS frameworks are elegant in that they derive metrics with clear biological interpretation that can be compared across populations and species (e.g., *z*, 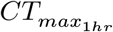). Here, we show that we can derive these same metrics directly from Equation 2.

As indicated above, we can model changes in the mean of any of the four parameters in Equation 2. Thermal sensitivity (*z*) and 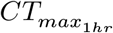 can be derived by modelling the effect of temperature on midpoints specifically (Figure 1b), as follows:

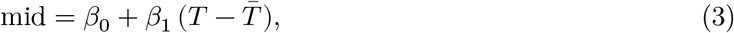

where mid is the log_10_-time at which survival equals the midpoint between the lower and upper asymptotes at assay temperature *T* (°C); *β*_0_ is the intercept giving the midpoint at the grand mean temperature 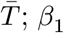 is the slope describing how the midpoint changes per °C increase in *T* ; and 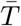 is the grand mean of assay temperatures used to mean-centre *T*, which decorrelates *β*_0_ from *β*_1_ and improves parameter estimation and interpretation (Schielzeth 2010).

When the experimental design estimates upper and lower limits of the response (i.e., low and up) near 0 and 1, the 4PL midpoint is equivalent to the classical *t*_50_. In this case, Equation 3 directly gives *z* = −1/*β*_1_, and the critical thermal limit at any reference time *t*_ref_ is 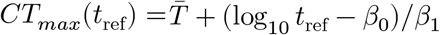 (with *t*_ref_ = 60 min for 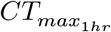).

More generally, the 4PL distinguishes between two related threshold definitions (Figure 1b). A *relative* threshold is defined by each curve’s own midpoint, *p* = (up +low)/2. On this scale, mid(*T*) is exactly the relative log_10_ *t*_50_, so *z*_rel_ = −1/*β*_1_ and *CT*_*max*,rel_ is read directly from Equation 3. An *absolute* threshold instead asks when survival reaches a fixed probability, such as 50% survival, regardless of the fitted asymptotes. When the low and up asymptotes differ from 0 and 1, this absolute threshold no longer coincides with the midpoint and requires an asymmetry correction (see Supplement Section 14):

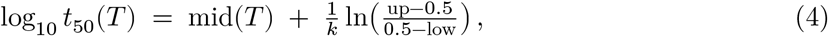

The corresponding absolute *CT*_*max*_ is obtained by applying the same correction on the temperature scale:

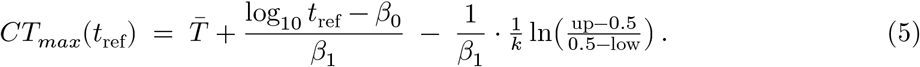

This distinction matters most when asymptotes or curve steepness vary with temperature. If the correction is constant across temperatures, absolute and relative *z* are the same, although *CT*_*max*_ may shift. If the correction changes with temperature, absolute log_10_ *t*_50_ is no longer parallel to the midpoint line, and *z*_abs_ can differ from *z*_rel_.

The same logic applies to any fixed mortality threshold. For example, log_10_ *t*_10_(*T*) is the time to 10% mortality (90% survival) and can identify the onset of population-level mortality, whereas log_10_ *t*_90_(*T*) is the time to 90% mortality (10% survival) and is closer to population collapse. Any mortality threshold can be estimated by using the corresponding probability in the correction term:

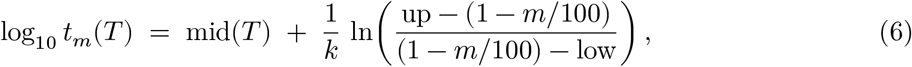

Here, *m* is the mortality percentage and 1 − *m*/100 is the corresponding survival probability used in the 4PL. Relative thresholds are useful when differences in asymptotes reflect baseline survival or incomplete response ranges. In contrast, absolute thresholds are useful when the biological question concerns a fixed survival or mortality probability. We therefore recommend reporting which threshold definition was used and, where asymptotes differ strongly among groups or temperatures, checking whether conclusions change under both definitions. In practice, there will often be little difference between relative and absolute summaries (i.e., *z* and *CT*_*max*_), but this distinction is important nonetheless.

Equivalently, the model can be reparameterised to estimate *z* and *CT*_*max*_ directly. Inverting the relations above (*β*_1_ = −1/*z* and 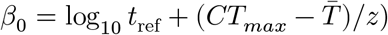 reparameterises the midpoint as mid(*T*) = log_10_ *t*_ref_ + (*CT*_*max*_(*t*_ref_) − *T*)/*z*, which substituted into Equation 2 gives

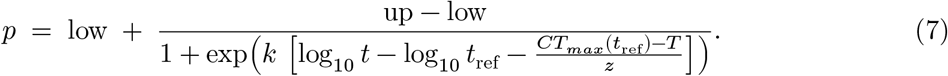

Now *z* (estimated on the log scale so it stays positive) and *CT*_*max*_ are estimated parameters in their own right, rather than quantities recovered from Equation 3 after fitting. Equation 7 uses the relative threshold (each curve’s own midpoint); an absolute threshold is obtained by adding the midpoint-scale asymmetry correction of Equation 4, 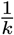 ln 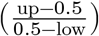, to mid(*T*) before solving for *CT*_*max*_. Again, temperature effects (and any interactions with moderators) should be included on low, up, and *k* to correctly estimate absolute thresholds, but are only needed on mid for relative thresholds.

A powerful aspect of the modelling approach described above is that uncertainty in all estimated quantities can be carried through the full analysis. The direct parameterisation (Equation 7) opens up ways of modelling *z* and *CT*_*max*_ that the two-stage approach cannot, while ensuring correct error propagation and inference. Each parameter in the model — `low`, `up`, `k`, `mid`, and, under Equation 7, *z* and *CT*_*max*_ — can be estimated separately for biologically meaningful strata (e.g., sex, population, species, treatment) by adding covariates and, where appropriate, random effects to the relevant parameter predictors (see Equation 11 below). This makes *a priori* tests of key hypotheses (e.g., do sexes or species differ significantly in *z* and *CT*_*max*_?) and variance decomposition (e.g., how much variation in *z* and *CT*_*max*_ arises across populations, species, or trials?) part of the model rather than a post-hoc comparison of fitted curves, all with correct inference. We fit the examples below in a Bayesian framework because posterior draws make uncertainty propagation straightforward, but the same likelihoods can also be fit in a frequentist framework, with uncertainty propagated by profiled likelihoods, the Delta method, or parametric bootstrap.

### Predicting heat injury and survival under dynamic thermal environments

A particularly powerful aspect of TDT/TLS models are their ability to translate dynamic temperature time series from nature to predict heat injury (HI) accumulation and thus population survival probability (Jørgensen *et al*. 2021; Li *et al*. 2023; Ørsted *et al*. 2022, 2024; Rezende *et al*. 2020). Heat injury accumulates as the time spent at a given temperature divided by the median lethal time at that temperature, integrated across the temperature trajectory *T* (*i*) as follows:

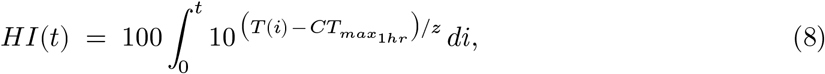

where *z* describes thermal sensitivity and 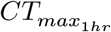 is the static temperature at which 50% of the population reaches the assayed endpoint after one hour of exposure. Larger *z* values indicate that a larger temperature increase is needed to reduce tolerance time by a factor of 10, so they correspond to lower sensitivity of tolerance time to warming. When *HI*(*t*) = 100%, the cumulative dose equals one *t*_50_ at the 1-hour reference, predicting 50% mortality (or 50% loss of the assayed sublethal endpoint, e.g. fertility; Ørsted *et al*. (2024)). Injury accumulates only when *T* (*i*) exceeds a critical temperature *T*_*crit*_ below which damage is repaired as fast as it accrues (Jørgensen *et al*. 2021; Ørsted *et al*. 2022).

A more natural prediction to make may instead be the population-level survival probability over time (Rezende *et al*. 2014, 2020). This has the advantage of allowing for a direct connection to population dynamical models which typically use survival, growth and fecundity as vital rates for population-level predictions (Noble *et al*. 2026). To predict survival, we can calculate the *effective exposure time* at a chosen reference temperature *T*_ref_. More effective exposure time is accumulated at higher temperatures and less at cool ones, in proportion to the scaling factor 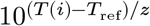,

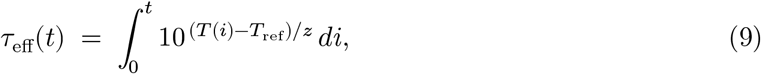

We can then evaluate the 4PL surface once at the cumulative effective exposure to obtain the predicted population-level survival fraction at every point in the trajectory:

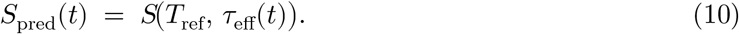

Here *S*(*T*, *τ*) is the survival surface fitted by the dose-response model, so *S*_pred_(*t*) is the survival probability for an organism held at *constant T*_ref_ after exposure duration *τ*_eff_(*t*). Direct error prop-agation of *z* and 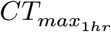 (using either Bayesian or frequentist approaches) allows one to easily model variation in outcomes, which can be helpful in making informed predictions in nature.

### Simulations - Evaluating statistical properties of the joint and two-stage modelling approaches

We assessed the performance of both the 4PL and two-stage approaches by estimating bias and 95% coverage for *z* and 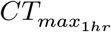 across simulation replicates. Bias was calculated as the estimated value minus the true value, whereas coverage was defined as the proportion of intervals containing the true parameter value. The 4PL approach estimated parameter uncertainty directly from the fitted model. In contrast, the two-stage approach required uncertainty to be propagated from the Stage-2 linear model using the Delta method, a first-order Taylor approximation for nonlinear functions of estimated parameters (Bolker 2008; Lappi & Luoranen 2018). For the two-stage estimates, we constructed intervals using both normal and *t*-distribution quantiles to evaluate the sensitivity of coverage to different assumptions about sampling uncertainty.

We tested the performance of the two-stage and 4PL approaches across scenarios designed to separate different sources of estimator bias and uncertainty. First, we simulated a strict-equivalence baseline in which survival spanned the full 0–1 range, there was no overdispersion, and only the dose-response midpoint varied with temperature. This scenario tested whether the two approaches recover the same quantities when the assumptions of the classical two-stage model are met. Second, we introduced beta-binomial overdispersion while keeping the same underlying dose-response shape, allowing us to isolate the cost of likelihood misspecification when binomial models are applied to overdispersed count data. Third, we simulated departures from the classical dose-response assumptions by allowing the upper asymptote to decline with temperature, both asymptotes to compress with temperature, or the curve steepness to change with temperature. These scenarios reflect empirical cases in which survival or sublethal responses do not span the full response range, or where the shape of the dose-response curve changes across assay temperatures (e.g., Bullard *et al*. 2026). We also ran sensitivity analyses that varied the strength of the upper-asymptote shift and the overall level of the upper asymptote.

Finally, we evaluated how different experimental designs impact estimates by simulating response curves from several static temperatures and exposure durations with modest numbers of individuals, samples, or replicate vials per treatment (Faber *et al*. 2024; Jørgensen *et al*. 2019; Li *et al*. 2023; Ørsted *et al*. 2024; Truebano *et al*. 2018). In the simulations, designs included 1, 3, or 5 replicate vials per temperature-by-time treatment, with 10 to 20 organisms per vial sampled from a uniform distribution. We also explored experimental designs that conducted experiments over short time spans across the different temperatures which we expected should impact model fit and therefore *z* and 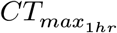 . For each scenario-by-design combination, we simulated 1,000 datasets and fitted both the two-stage (both relative and absolute summaries) and 4PL approaches. Overall, this amounted to 29,000 simulations across the 29 scenario-by-design combinations. We report Monte Carlo standard errors (MCSE) for the bias and coverage summaries to better understand across simulation uncertainty (DiRenzo *et al*. 2023; Koehler *et al*. 2009; Williams *et al*. 2024).

#### Simulation results

When the data-generating process matched the classical assumptions, with survival spanning the full 0–1 range and temperature affecting only the dose-response midpoint, the two-stage approach performed similarly to the joint 4PL model, producing little bias and near-nominal coverage when confidence intervals were constructed with *t*-distribution quantiles. Using normal (z) quantiles instead produced anti-conservative intervals with coverage well below the nominal 95% level (Supplement 10.1 and 10.2).

In contrast, the two-stage approach resulted in biased *z* and 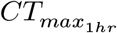, often with poor coverage, when temperature affected the upper asymptote (`up`) or the rate of decline (`k`) (Figure 2). Bias for both *z* and 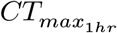 from the two-stage approaches was particularly severe as the upper survival threshold differed from 1, or decreased with temperature (Figure 3). These biases were consistent even with larger sample sizes (Supplement 10.6), but they became most consequential under exactly the types of experimental designs used in practice: sparse temperature-duration grids, short exposure windows, and low replication (Figure 4). The joint 4PL remained near-unbiased and well-calibrated across these designs, whereas two-stage bias grew and interval coverage collapsed as grids became sparser, replication reduced, or exposure windows shortened — most severely at the shortest exposure cap, where two-stage coverage of *z* fell below 0.05 even with the *t*-correction (Figure 4; full per-cell detail in Supplement 10.6).

**Figure 2:**
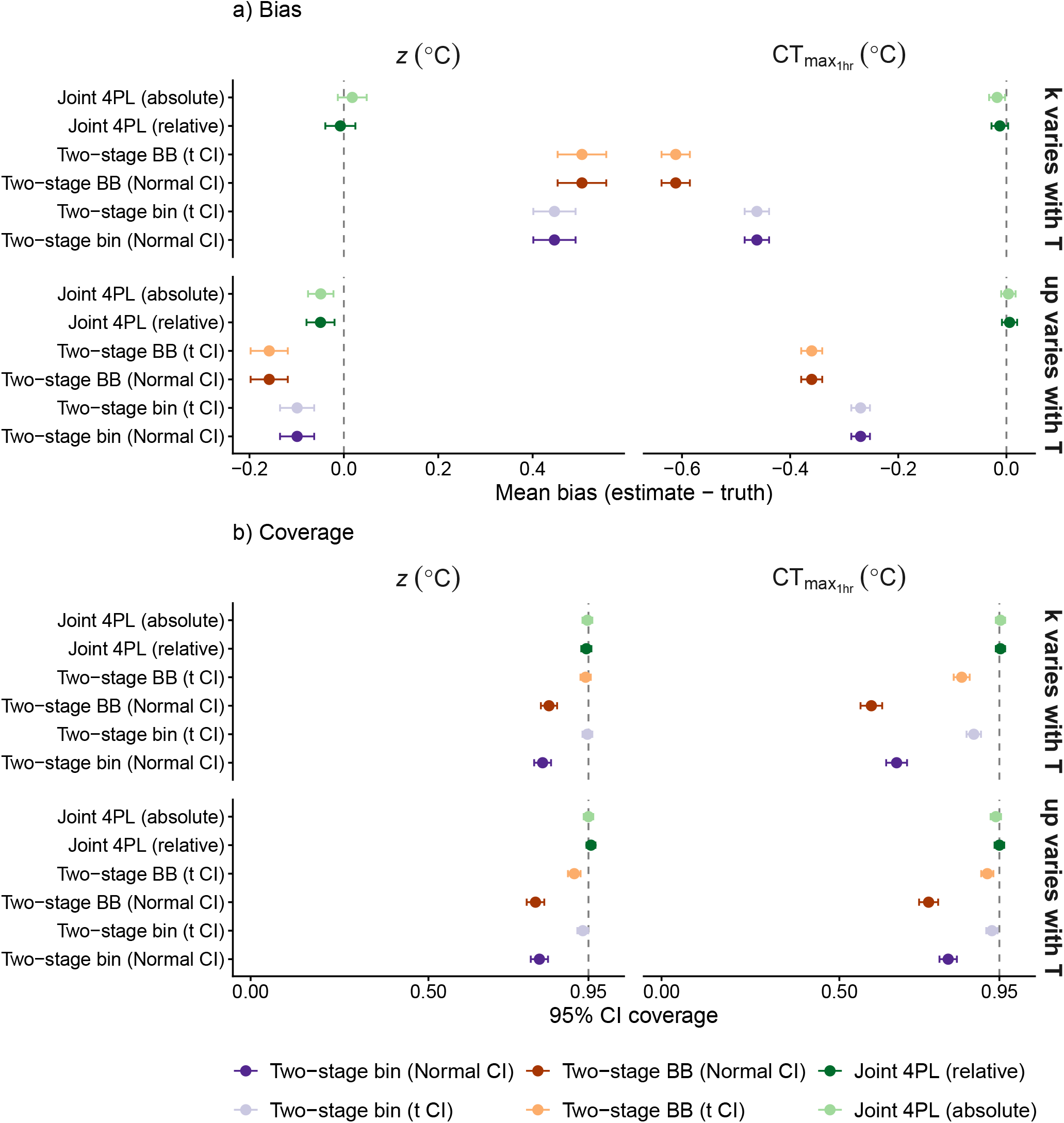
Simulation performance when the data-generating process violates classical TDT shape assumptions. Row labelled ‘up varies with T’: upper asymptote up declines 0.01 per °C from 0.92 at *T* = 34 °C, with low = 0.05 and *k* = 8 constant in *T* . Row labelled ‘k varies with T’: slope *k* ranges from 7 at 30 °C to 9 at 38 °C, with up = 0.92 and low = 0.05 constant. Both rows use 5 replicate vials per (T × duration) cell, *N*_*sim*_ = 1000 simulated datasets per row, and beta-binomial overdispersion (*ϕ* = 5). Panel a): mean signed bias in *z* (left column) and 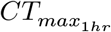 (right column) with 95% Monte Carlo uncertainty intervals (mean ± 1.96 MCSE). Panel b): 95% interval coverage with the same column layout. Dashed lines mark zero bias and nominal coverage. Two-stage Normal- and t-quantile variants share point estimates and differ only in Stage-2 delta-method interval construction. 4PL = four-parameter log-logistic, BB = beta-binomial, bin = binomial, CI

**Figure 3:**
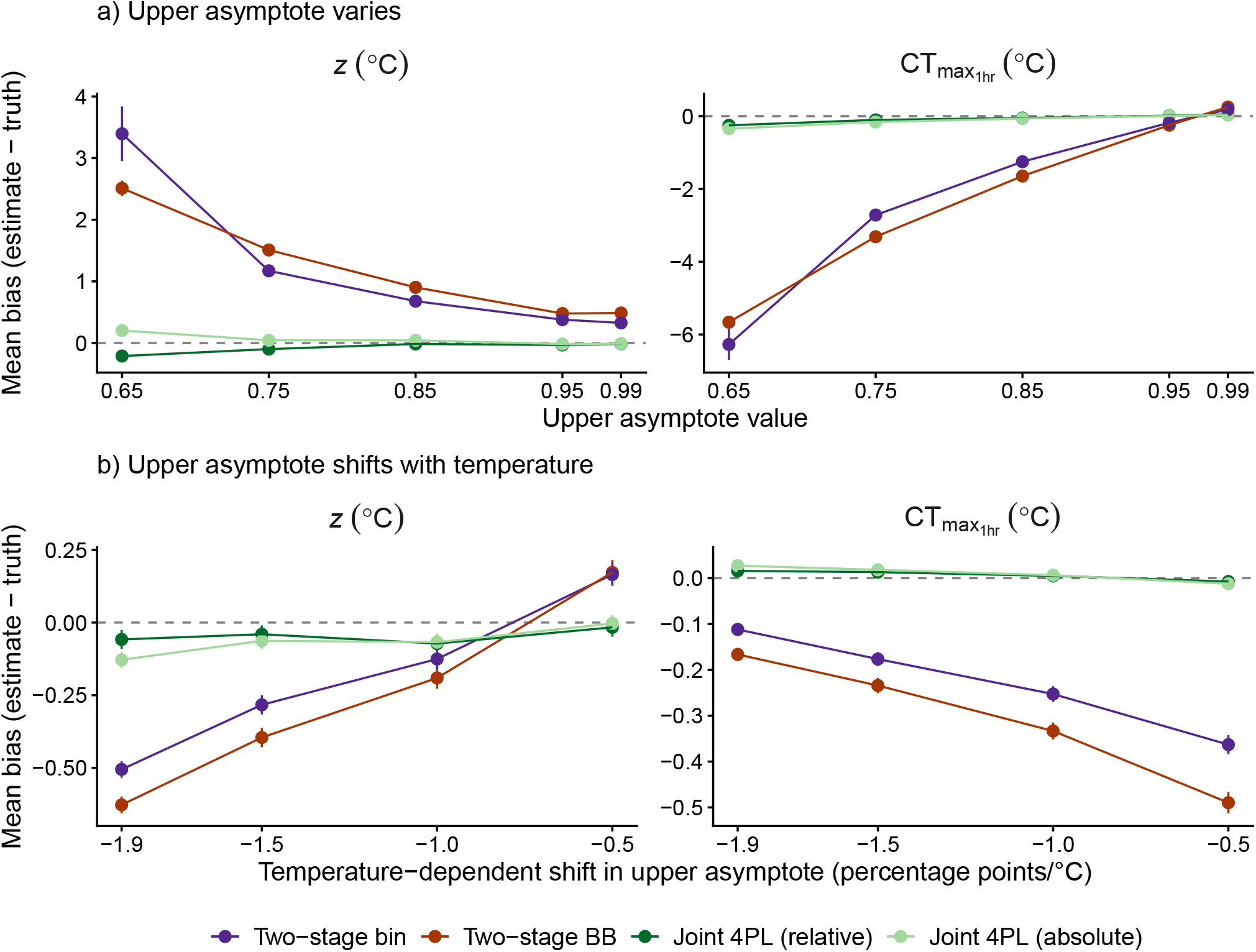
Bias in *z* and 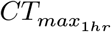 under two departures from a full upper asymptote. Panel a) (upper asymptote varies): *β*_*up*_ = 0 (no temperature-dependent shift), but the upper asymptote is held at a reduced level across all temperatures, so the maximum attainable survival is lower throughout the grid. Panel b) (upper asymptote shifts with temperature): the upper asymptote at *T* = *T* is fixed at up_0_ = 0.92 while its per-°C shift (*β*_*up*_) is varied, so up stays high at cool assays and declines progressively at hotter ones. Both panels: low = 0.05, *k* = 8, mid linear in *T* with slope −0.15, beta-binomial likelihood (*ϕ* = 5), *n*_*reps*_ = 5, *N*_*sim*_ = 1000 datasets per simulated scenario. Points are mean signed bias with 95% Monte Carlo uncertainty intervals (mean ± 1.96 MCSE); dashed lines mark zero bias. Left column: *z*; right column: 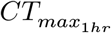 . Only the four distinct point-estimate methods are shown: the two-stage Normal- and t-quantile variants share point estimates within each Stage-1 family, so bias is identical between them. Coverage differences are presented in the supplement for panel a, Supplement 10.5 and panel b, Supplement 10.4.

**Figure 4:**
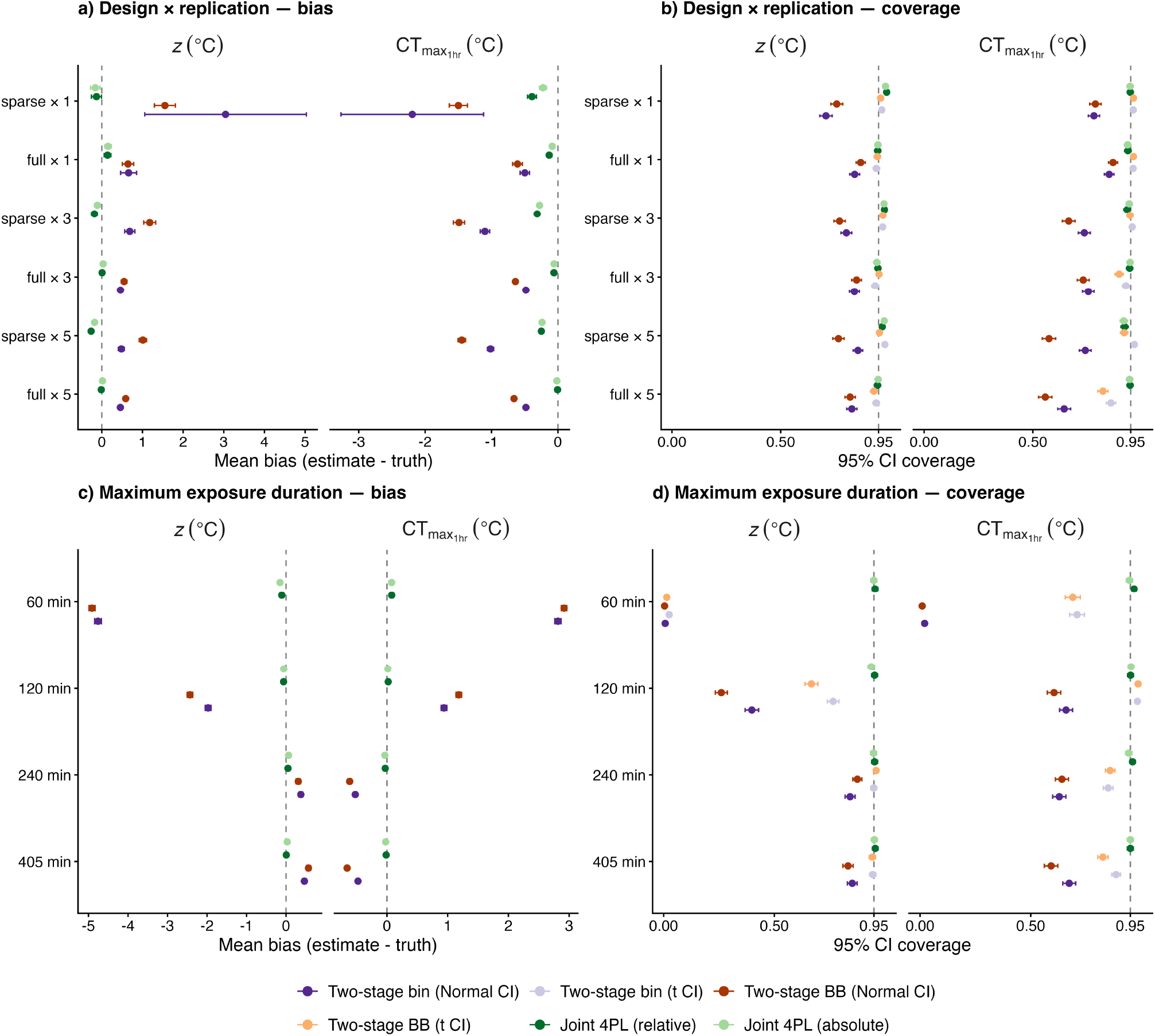
Estimator performance across experimental designs. Panels a) and b) vary experimental design and replication: a full (5 temperatures × 6 durations) versus sparse (3 × 4) temperature– duration grid, crossed with 1, 3, or 5 replicate vials per treatment combination. Panels c) and d) vary the maximum exposure duration (the longest assay window: 60, 120, 240, or 405 min) on the full grid with five replicates. Panels a) and c) show mean signed bias and panels b) and d) show 95% interval coverage for thermal sensitivity *z* (left) and 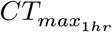 (right). Points are Monte Carlo means with 95% Monte Carlo uncertainty intervals (mean ± 1.96 MCSE); dashed lines mark zero bias and nominal 0.95 coverage. Bias panels show the four distinct point estimators (the two-stage Normal- and *t*-CI variants share point estimates); coverage panels additionally distinguish the *t*-CI variants. All simulated scenarios use a beta-binomial data-generating process (*ϕ* = 5) with a fixed 4PL shape (up = 0.92, low = 0.05, *k* = 8, midpoint linear in *T*); *N*_*sim*_ = 1000 datasets per simulated scenario. Full scenario detail and RMSE are in Supplement 10.6.

### Worked examples

We applied the 4PL modelling framework to four case studies spanning lethal counts, continuous-proportion physiological data and sublethal responses using four case studies: 1) three cereal-aphid species (Li *et al*. 2023), 2) snow gum leaf PSII function (Arnold *et al*. 2026), 3) zebrafish larvae under varying oxygen levels (Saruhashi *et al*. 2026) and 4) vinegar fly mortality, heat coma and productivity (Ørsted *et al*. 2024). These examples show how the same fitted response surface can estimate *z, CT*_*max*_ and, for lethal endpoints, *T*_*crit*_, compare groups, and propagate uncertainty into heat-injury and survival predictions. The workflow is implemented in `bayesTLS`; the online supplement provides installation instructions, a tutorial and the full model specifications (Supplement Sections 1–4).

For the main grouped case studies, we used the direct parameterisation in Equation 11, placing species, oxygen treatment or recovery condition directly on *CT*_*max*_ and *z*. The vinegar-fly sex comparison was fit in the equivalent midpoint form for demonstration, with `sex` and its temperature interaction on mid. Count responses were fit with beta-binomial likelihoods, while proportions in the snow gum example were fit with a beta likelihood. We fitted all models using a Bayesian implementation, with `brms` (v2.23.0; Bürkner (2017)) and Stan (v2.38.0; Carpenter *et al*. (2017)) `via bayesTLS` (v1.0.0). Equivalent frequentist models are available via `freqTLS`, and we demonstrate their equivalence in Supplement Section 13.

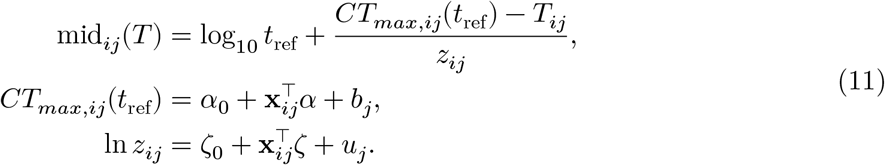

For the lethal aphid and vinegar-fly examples, we also used posterior draws to predict heat injury and survival under field or field-derived temperature traces, with and without an illustrative Sharpe– Schoolfield temperature-dependent repair function (*sensu* Arnold *et al*. 2025). Full formulas, priors, sampler controls, diagnostics and temperature-trace details are reported in Supplement Section 4. The survival estimates use posterior draws for *z*, 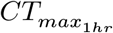, and *T*_*crit*_ (estimated as the temperature at which injury starts to accumulate), linking laboratory tolerance to predictions of survival in the field.

### Cereal aphids (*Metopolophium dirhodum, Sitobion avenae, Rhopalosiphum padi*)

Li *et al*. (2023) found pronounced differences in heat tolerance and sensitivity among species, which likely explain changes in cereal-aphid communities. Our 4PL model recovers similar findings, with *R. padi* having the highest 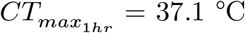 (exceeding *M. dirhodum* by 2.0 [1.8, 2.2] °C, *p*_MCMC_ < .001) and *M. dirhodum* the shallowest thermal-death-time slope (largest *z* = 4.8 °C) and the lowest 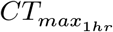 (Figure 5a). However, because the 4PL is fit to the raw survival counts, *R*^2^ was more realistic (0.695, 95% CrI: 0.671-0.715), and well below the reported *R*^2^ values from the two-stage regressions (0.97-0.996; Supplement Section 6.2).

**Figure 5:**
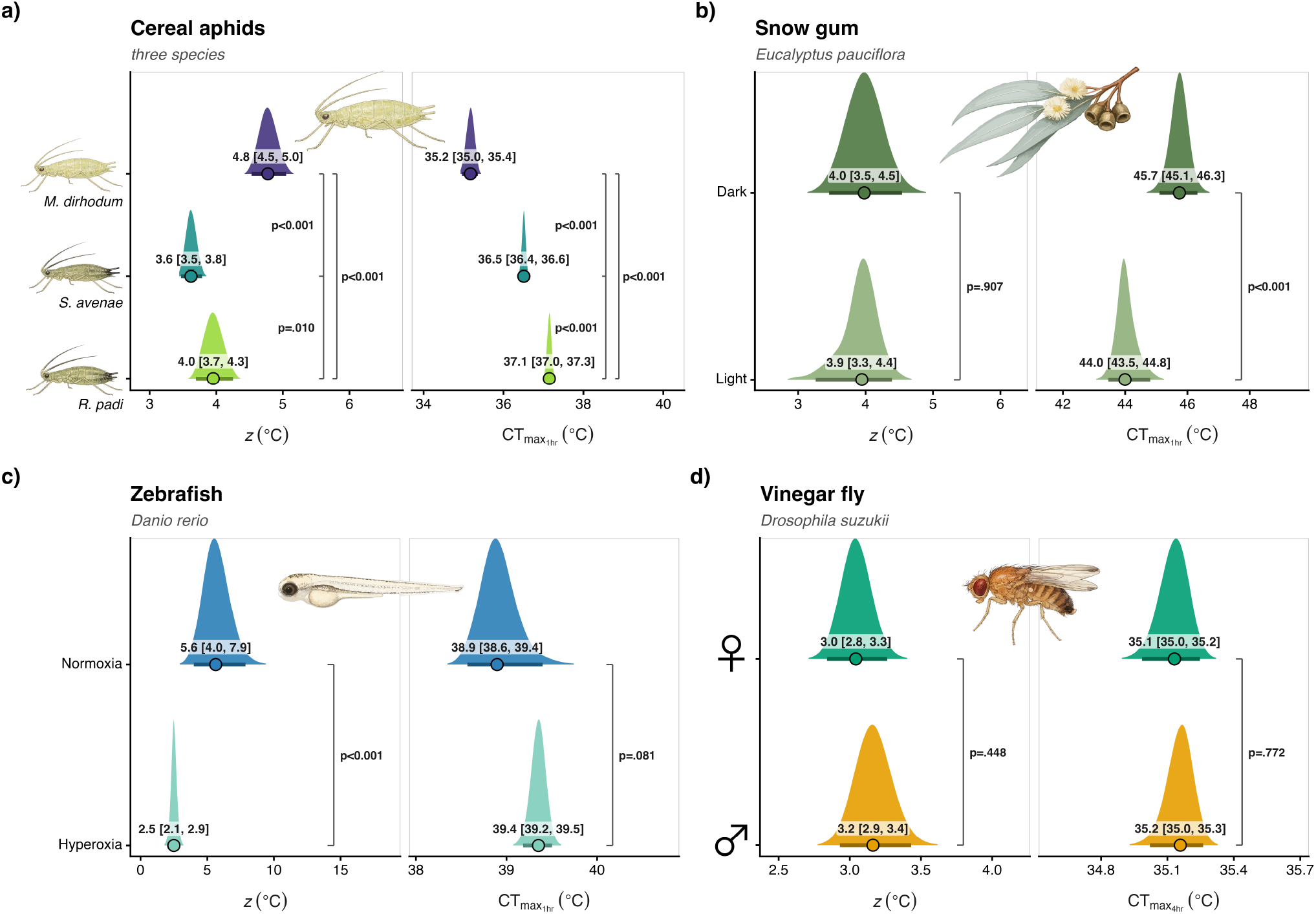
Thermal sensitivity (*z*, left of each panel) and the critical thermal limit (*CT*_*max*_, right) estimated by the joint Bayesian four-parameter log-logistic (4PL) model across four case studies: a) three cereal-aphid species (*Metopolophium dirhodum, Sitobion avenae, Rhopalosiphum padi*; Li *et al*. (2023)), b) snow gum (*Eucalyptus pauciflora*) leaf photosystem II function under dark vs 90-min light treatment during recovery from heat stress (Arnold *et al*. 2026), c) zebrafish (*Danio rerio*) larvae under normoxia and hyperoxia conditions (Saruhashi *et al*. 2026), and d) vinegar fly (*Drosophila suzukii*) males and females (Ørsted *et al*. 2024). Larger *z* values indicate lower sensitivity of tolerance time to warming (i.e., a larger increase in temperature is required to reduce tolerance time by a factor of 10). Filled ridges are posterior densities, coloured by group. Points and error bars correspond to posterior medians and 95% credible intervals, which are also printed beside each distribution. The *CT*_*max*_ is predicted for an exposure duration of 1 hour for all case studies except vinegar flies, where we used 4 hours as per the original case study. Within each panel, grouping variables were fit within a single joint model, so that differences among groups are evaluated as pairwise posterior contrasts, annotated as brackets and significance (alpha = 0.05) labelled with *p*.

Using field temperatures from the same site (Wuhan, China), we also show how the 4PL can predict field mortality with uncertainty propagated from that same model: survival without repair is predicted to fall to roughly 2.0-2.1% by late June, while physiological repair raised end-of-June survival by 4.8 [0.3, 24.0]% for *R. padi* and 1.4 [0.3, 5.1]% for *S. avenae*, but barely changed it (< 0.1%) for the least-tolerant *M. dirhodum* (Figure 6d-f). The model therefore moves inference from ranking laboratory tolerance to predicting which species is most likely to experience mortality during natural fluctuating temperature regimes.

**Figure 6:**
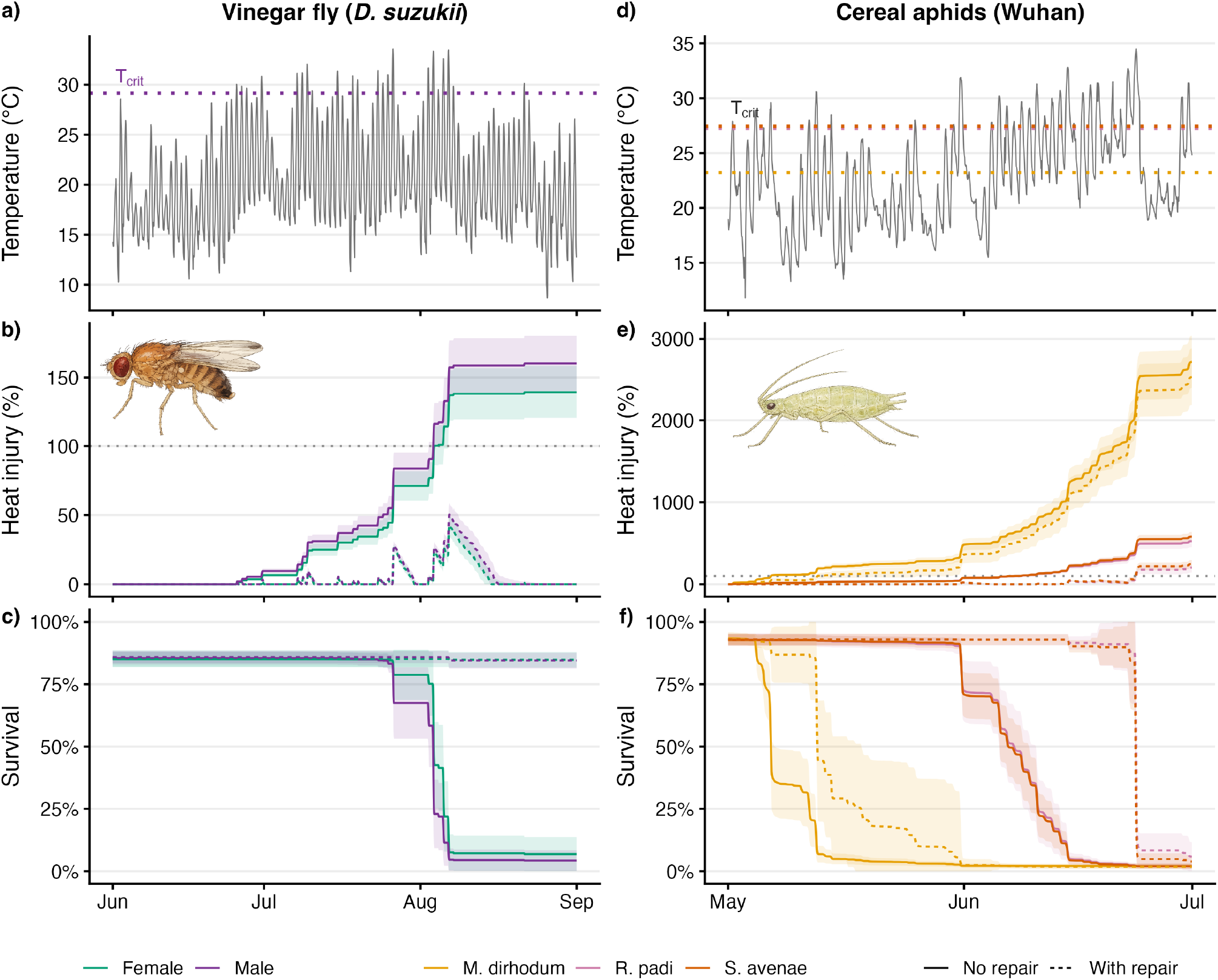
Heat-injury accumulation and predicted survival under field temperature-time series for the vinegar fly (*Drosophila suzukii*, by sex; panels a-c) and three cereal aphids (*Metopolophium dirhodum, Sitobion avenae, Rhopalosiphum padi*; panels d-f). (a, d) Temperature-time series: NicheMapR (Kearney & Porter 2017) microclimate temperatures in the shade for the vinegar fly (Rennes, France, Jun–Aug 2018; Ørsted *et al*. (2024)) and hourly air temperatures for the aphid site (Wuhan, May–June 2016; Li *et al*. (2023) site and period, re-sourced from the Open-Meteo ERA5 reanalysis), each with the critical temperature threshold *T*_*crit*_ (dotted line). (b, e) Cumulative heat injury; 100% corresponds to one *LT*_50_ dose, i.e. the thermal dose predicted to cause a 50% decline in survival (dotted grey line). (c, f) Predicted survival. Both case studies show predictions without (solid line) or with (dashed line) physiological repair at permissive temperatures. Fly panels are coloured by sex, aphid panels by species. Bands are ±1 SE across posterior draws. Repair uses an illustrative Sharpe–Schoolfield kernel with *r*_*ref*_ = 0.005 LT50-dose h^−1^ at the reference temperature; this repair function is illustrative and was not derived empirically.

### Snow gum (*Eucalyptus pauciflora*)

Arnold *et al*. (2026) showed that light conditions alter estimates of heat damage to photosystem II (PSII), an ecologically important endpoint because the leaves of snow gum (and most other canopy-forming plants) experience the most extreme heat during daylight, often with high solar input. By fitting a 4PL model to continuous PSII proportions (post/pre *F*_*v*_/*F*_*m*_) with a beta distribution, we reproduce the key result that light conditions following heat stress lower heat tolerance, while clarifying the mechanism at play: light treatment reduced 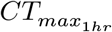 by 1.7 [1.0, 2.2] °C (*p*_MCMC_ < .001), but did not affect *z* (*p*_MCMC_ = .91; Figure 5b) relative to the dark treatment. Therefore, light mainly lowers the thermal load required to impair PSII, but not the rate at which such load accumulates over time, implying that daylight heatwaves may push leaves into heat stress at lower temperatures than dark-recovery assays suggest.

### Zebrafish (*Danio rerio*)

Saruhashi *et al*. (2026) found that oxygen modulates heat tolerance in zebrafish larvae, an important result because freshwater heatwaves often coincide with changing oxygen availability. Despite the sparse design, which would make it challenging to use a two-stage analysis to estimate *CT*_*max*_ and *z*, we show that the 4PL can still be used. Our model recovers higher heat tolerance under hyperoxia (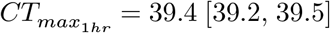 vs 38.9 [38.6, 39.4] °C under normoxia; Figure 5c), but also shows that hyperoxia *steepens* the TDT relationship relative to normoxia (Δ*z* = -3.1 [-5.4, -1.5] °C, *p*_MCMC_ < .001). The 4PL therefore provides stronger ecological inference by demonstrating that oxygen does not simply raise heat tolerance, it also changes how fast injury is predicted to accumulate during heat stress, so its benefit depends on exposure duration as well as temperature intensity.

### Vinegar fly (*Drosophila suzukii*)

Ørsted *et al*. (2024) found that heat coma and declines in productivity occur at temperatures below survival limits but follow traditional TDT relationships, implying that sublethal effects have large implications for population dynamics. Our re-analysis of their survival data recovers similar variation in thermal tolerance and sensitivity between sexes (*z* = 3.0 [2.8, 3.3] °C for females and 3.2 [2.9, 3.4] °C for males; 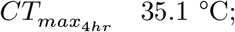 35.1 °C Figure 5d), but estimates the sex contrast directly in a unified model and finds no detectable difference (Δ*z* = 0.1 [-0.2, 0.5] °C, *p*_MCMC_ = .45; *CT*_*max*_ *p*_MCMC_ = .77). As with the aphids, retaining the raw-response variance gives more realistic fit summaries (Bayesian *R*^2^ = 0.90 for females and 0.79 for males; Supplement Table S18). In the supplement, we also extend modelling approach to heat coma (Supplement Section 9.0.0.0.1) and productivity (Supplement Section 9.0.0.0.2), including a hurdle model that separates reproductive failure from changes in offspring number among reproducing flies.

We generated predictions on heat injury and predicted mortality using a microclimate temperature-time series for Rennes, France. Such modelling illustrates how patterns of population mortality for both sexes can be predicted in nature with and without repair (Figure 6a-c), showing how repair (even at small rates) can have important population-level consequences.

## Discussion

In this study, we show that conventional TDT/TLS analyses can be reparameterised using a four-parameter log-logistic model (4PL) to offer more robust, accurate, and generalisable predictions of the impacts of extreme heat on organisms. The 4PL model provides a flexible and statistically rigorous framework for estimating thermal tolerance and thermal sensitivity, recovering familiar quantities such as *z* and *CT*_*max*_. At the same time, the model derives estimates from the full temperature-duration response surface and propagates uncertainty into every derived quantity. Simulations show that the joint model reproduces classical estimates when two-stage assumptions hold, but is more reliable when experiments are sparse, replication is low, exposure windows are short, or the response bounds (`low, up`) depart from 0 and 1 (Figure 2–Figure 4). Using four case studies across diverse plant and animal taxa, we show how the 4PL model recovers biologically plausible TLS estimates, with *z* values falling within the range typically reported for thermal tolerance landscapes (approximately 2-7 °C; Rezende *et al*. (2014)), and unlocks opportunities to extend inference to sublethal endpoints, group contrasts, and survival under fluctuating temperature regimes.

One of the key challenges in global change biology is to understand the ecological and evolutionary drivers of thermal sensitivity and tolerance across life (Jørgensen *et al*. 2022; Rezende *et al*. 2014). Species, populations, life stages, and organisms exposed to various experimental treatments are likely to differ in the temperature at which damage starts to accumulate (*T*_*crit*_), the rate at which damage accumulates (*z*), and how much heat load they can endure (*CT*_*max*_) (Arnold *et al*. 2025; Li *et al*. 2023; Ørsted *et al*. 2024; Saruhashi *et al*. 2026; Youngblood *et al*. 2025). We need robust statistical approaches that allow us to quantify differences in how temperatures impact biological function, which was not previously possible with conventional two-stage analysis pipelines. Using broad case studies, we illustrate the power of the 4PL modelling approach in estimating these parameters and how they differ among groups of particular interest in global change biology. In cereal aphids, our model was able to statistically compare differences in thermal tolerance and sensitivity among species, and to link interspecific differences in heat tolerance measured in the laboratory to predicted survival under natural temperature regimes. In the snow gum, our model was able to statistically separate how light conditions affect the thermal load required to impair PSII and the sensitivity of that response, while accounting for their covariance. In zebrafish, we show that oxygen availability changes the entire shape of the thermal tolerance surface, not simply heat tolerance. In vinegar flies, we show that the sexes respond similarly to heat stress, but that vital rates (i.e., mortality, coma, productivity) follow distinct thermal tolerance landscapes and fail at different temperatures under extreme heat. These ecological inferences provide new insights into the mechanisms that may underlie vulnerability to extreme heat, not only alternative ways for estimating *z* or *CT*_*max*_.

Because the 4PL model fits raw response data in a single hierarchical analysis, it also creates new opportunities for asking where variation in responses comes from. Clones, individuals, tissues, sexes, populations, or treatments can be included in the same model, on any of the 4PL parameters, to estimate genetic, individual, tissue-specific, and environmental sources of variance in *z* and *CT*_*max*_. The posterior distributions of the underlying 4PL parameters (`low, up, k`, and `mid`) can also be contrasted to determine whether groups differ in baseline survival, maximum attainable survival, curve steepness or thermal sensitivity. This kind of variance decomposition is not possible with a two-stage pipeline because the first stage collapses the response surface to a small set of point estimates, and the second stage treats those estimates as data. Full propagation of sampling uncertainty should also make tolerance and sensitivity estimates more useful, and less biased, for robust meta-analyses, comparative studies, and systems models of biological responses to extreme heat (Noble *et al*. 2026; Pottier *et al*. 2024).

The same modelling framework extends beyond lethal count data to many sublethal measures that bbare of interest to ecologists and evolutionary biologists and could arguably be more important in assessing vulnerability to environmental change (Arnold *et al*. 2026; Bretman *et al*. 2024; Ørsted *et al*. 2026; Snook *et al*. 2026). In the supplement, we show how derivatives of the 4PL model can be used for time-to-event responses, mixture models that separate response and non-response components, Gamma-distributed responses, and continuous proportions. This modelling framework can also be expanded to include interacting stressors (e.g., oxygen, acidification, pollutants), as well as temperature-dependent effects on acclimation, physiological repair, and behaviour, which can all modulate the accumulation of heat injury under fluctuating temperature regimes (Arnold *et al*. 2025). More generally, any dose-response process in which biological responses depend jointly on stress intensity and exposure duration could be analysed with this framework, including toxicology studies where toxicant concentration replaces temperature. Therefore, the 4PL modelling framework opens avenues to build greater consistency in reporting and analytical choices across fields, which can have consequences for ecological inference (Gould *et al*. 2025).

While the 4PL framework is powerful, it also has limitations. Fitting models with temperature-dependent effects on all parameters may cause some to be estimated poorly in very small datasets. However, our simulations show that, even with small sample sizes and less rigorous designs, the 4PL can recover correctly calibrated estimates, which offers opportunities to apply this framework to taxa where large sample sizes are difficult to achieve. Nonetheless, models may at times need to be simplified, for example by ignoring temperature-dependent shifts in `up`, `low`, or `k` if they are not well supported by the data. Bayesian implementations also require priors that are appropriate for the biological system, although `bayesTLS` is designed to make these models more accessible to experimental and comparative biologists. Similar models can also be fitted in a likelihood framework, with uncertainty propagated using profiled likelihood, parametric bootstrapping or the Delta method (Bolker 2008). We provide this option through `freqTLS`. However, propagating uncertainty into survival and heat-injury predictions is more straightforward within the Bayesian framework, which also incorporates prior information. A major next step is to estimate repair parameters directly from experiments (Arnold *et al*. 2025; Buckley *et al*. 2025; Ørsted *et al*. 2026), because our illustrative Sharpe–Schoolfield repair model (*sensu* Arnold *et al*. (2025)) shows that predictions of mortality can change substantially when repair is included. Future models that combine heat injury, repair, acclimation, behaviour, and microclimate validation will be essential for translating laboratory tolerance surfaces into robust forecasts of survival in nature (Arnold *et al*. 2026; Faber *et al*. 2026; Noble *et al*. 2026).

Taken together, the 4PL modelling framework allows more flexible, robust, and generalisable inferences of thermal tolerance, sensitivity, and vulnerability of organisms to extreme heat. This framework not only helps us better understand the ecological and evolutionary drivers of biological responses within and among species, but also helps translate laboratory estimates into uncertainty-aware forecasts of survival under fluctuating temperature regimes in the field. Therefore, this modelling framework is likely to improve our ability to predict organisms’ vulnerability to increasingly intense, frequent, and long-lasting extreme heat events that will be common in the future.

## Acknowledgements

We thank Johannes Overgaard, Joey Bernhardt, Wilco Verbeck, Jennifer Sunday, Ignacio Peralta-Maraver, Arnaud Sentis and Jeff Arnoldi for discussions during an extreme heat workshop supported by the ClimateCountDown ERC Consolidator Grant, funded by the European Research Council under the European Union’s Horizon 2020 research and innovation programme (Grant agreement no. 101169909). We would also like to thank Gwenaëlle Deconninck, Charlie Cornwallis, and Tobias Uller for helpful comments. D.W.A.N was supported by an ARC Future Fellowship (FT220100276) and thanks the ANU, Lund University, the ANU Hansen Scandinavian Friendship Grant, and the Royal Physiographic Society of Lund for funding that supported the sabbatical during which this paper was written. P.P. was supported by a Wenner-Gren Stiftelserna postdoctoral fellowship (UPD2024-0239). P.A.A. is supported by an ARC Discovery Project (DP240100177). S.N. was supported by a Canada Excellence Research Chair (CERC-2022-00074).

## Author contributions

D.W.A.N.: Conceptualization, Methodology, Software, Data Curation, Formal Analysis, Visualization, Writing – Original Draft. P.P.: Conceptualization, Methodology, Software, Data Curation, Formal Analysis, Visualization, Writing – Original Draft. P.A.A.: Conceptualization, Methodology, Investigation, Resources, Writing – Review & Editing. S.N.: Conceptualization, Methodology, Software, Investigation, Resources, Writing – Review & Editing. All authors contributed to interpretation of results, reviewed and edited the manuscript, approved the final version, and agree to be accountable for the work.

## Data and code availability

Analysis scripts, case-study data, committed model fits, and rendered outputs are available at https://github.com/daniel1noble/bayesTLS. Cached simulation outputs and model fits are archived on OSF (https://osf.io/c6dxy) and can be fetched with make data; the raw simulation files are regenerable from the simulation scripts in the repository. The online supplement, tutorial, and bayesTLS installation instructions are available at https://daniel1noble.github.io/bayesTLS/. We also provide a sister package, freqTLS, which develops these models in a likelihood framework.

## AI Declaration

During the preparation of this manuscript, the authors used generative AI tools, including OpenAI ChatGPT/Codex (vers. 5.5) and Claude Code (vers. opus 4.8), to assist with language editing, structural refinement, mathematical derivations, code building and checking, R package construction and formatting. All scientific ideas, analyses, interpretations, and conclusions were developed and verified by the authors. Any AI-assisted output was critically reviewed and edited by the authors, who accept full responsibility for the final content of the manuscript, supplement, code, and associated R packages.

